# PlantRNA 2.0: an updated database dedicated to tRNAs of photosynthetic eukaryotes

**DOI:** 10.1101/2021.12.21.473619

**Authors:** Valérie Cognat, Gael Pawlak, David Pflieger, Laurence Drouard

## Abstract

PlantRNA (http://plantrna.ibmp.cnrs.fr/) is a comprehensive database of transfer RNA (tRNA) gene sequences retrieved from fully annotated nuclear, plastidial and mitochondrial genomes of photosynthetic organisms. In the first release (PlantRNA 1.0), tRNA genes from 11 organisms were annotated. In this second version, the annotation was implemented to 51 photosynthetic species covering the whole phylogenetic tree of photosynthetic organisms, from the most basal group of Archeplastida, the glaucophyte *Cyanophora paradoxa*, to various land plants. Transfer RNA genes from lower photosynthetic organisms such as streptophyte algae or lycophytes as well as extremophile photosynthetic species such as *Eutrema parvulum* were incorporated in the database. As a whole, circa 37 000 tRNA genes were accurately annotated. In the frame of the tRNA genes annotation from the genome of the Rhodophyte *Chondrus crispus*, non-canonical splicing sites in the D- or T- regions of tRNA molecules were identified and experimentally validated. As for PlantRNA 1.0, comprehensive biological information including 5’- and 3’-flanking sequences, A and B box sequences, region of transcription initiation and poly(T) transcription termination stretches, tRNA intron sequences and tRNA mitochondrial import are included.

## INTRODUCTION

Beyond their major role in translation as carriers of amino acids, transfer RNAs (tRNAs) have many non-canonical functions and are involved in a multitude of biological processes, as already emphasized in PlantRNA 1.0 database (1) or in more recent reviews (e.g.(2) (3)). For instance, the key role played by tRNAs in the production of tRNA-derived small RNAs (tDRs) (4) involved in diverse functions is expanding rapidly (5). Furthermore, there is growing evidence that not only tRNAs but also tRNA genes appear to play important roles. In particular, tRNA genes can be involved in the organization of the nuclear genome and affect the chromatin structure (e.g. (6) (7) (8) (9)). These genes are either scattered or clustered throughout the genomes (10) (11). Their number greatly varies between the nuclear genomes of eukaryotes (8), and likely their genomic distribution varies as well but this remains poorly explored. To better decipher how tRNA genes contribute to genome evolution it is therefore essential to retrieve complete sets of tRNA genes with the highest possible accuracy. Photosynthetic organisms, with the presence of three genomes (*i.e*. nuclear, plastidial, and mitochondrial) represent the most complex model of eukaryotic cells to visualize the evolution of the tRNA genes population in terms of gene content, organization and expression. In 2011, we thus decided to develop a tRNA database dedicated to photosynthetic organisms and the first version of PlantRNA (12) has provided information on 11 eukaryotes of the photosynthetic lineage. Although covering evolutionary distinct branches of the photosynthetic lineage, this database was rather limited. During the last decade, the number of completed nuclear, plastidial and mitochondrial genome sequences has considerably increased. This recent accumulation of knowledge has now made it possible to include the tRNA gene sequences of about 40 additional species representative of the phylogenetic tree of the Archeplastida lineage. This includes species representative of terrestrialisation, vascularisation, flowers emergence, or growth in extreme environments as well as two rhodophytes whose genomes contain permuted tRNA genes or peculiar intronic sites. The genomes of a cryptophyte, a haptophyte and two heterokontophytes were also annotated. The recently optimized version of tRNAscan-SE program (13) was used to predict and annotate tRNA genes as accurately as possible. Nevertheless, manual curation was still essential at different levels, for example, to avoid false annotations due to contamination from external (*e.g*. bacterial) or organellar DNA during nuclear genome sequencing programs, to eliminate remaining tRNA pseudogenes that escape detection, or to identify missing plastidial tRNA genes containing group II introns. Biological information relevant to tRNA biology (*e.g*. intron sequences, flanking sequences controlling tRNA gene expression, tRNA mitochondrial import) is also made available.

As a whole, *circa* 37 000 novel tRNA genes were annotated, thus providing the largest curated tRNA database of photosynthetic organisms.

## DATABASE CONTENT AND WEB INTERFACE

The current release incorporates data sources of the previous version for 7 species (12), novel data sources for 4 updated genomes, and a choice of 40 new data sources selected on the basis of the interest of the species regarding their position in the phylogenetic tree of the photosynthetic lineage. For instance, they cover 7 genomes of streptophyte algae belonging to the lineage being at the transition step from aquatic algae to land plants, the genomes of a lycophyte (*Selaginella moellendorffii*) the first non-seed vascular plant genome reported (14), a fern (Azolla filiculoides), one of the fastest-growing plants, or *Amborella trichopoda*, representative of the sister lineage of all flowering plants. Plant species were also chosen because of their growth in peculiar environments such as the cave plant *Primulina huajiensis* (15) or the halophytic plant *Eutrema parvulum* (also called *Schrenkiella parvula* or *Thellungiella parvula*) (16,17). The quality of the genome annotations was also taken into account whenever possible. In addition, when available, plastidial and mitochondrial genomes were also used as sources. Whole nuclear, plastidial, and mitochondrial genomes were scanned by tRNAscan-SE 2.0 (13) and automatically processed by a homemade workflow (available at https://github.com/ibmp-bip/tRNAflow) and then manually curated and annotated as in (18). Figure 1 shows the phylogenetic tree rof the photosynthetic organisms from which the tRNA genes were retrieved. The sources of genome sequences listed in Supplemental Table S1 are accessible via the entry point “Species” on the PlantRNA website. As in version 1.0 (12), additional data including linear secondary structure, A and B boxes, flanking sequences controlling tRNA gene expression, intronic sequences, mitochondrial tRNA import either experimentally proven or predicted, are indicated. Regarding intronic sequences, introns predicted at non-canonical positions in the red alga, *Chondrus crispus*, were experimentally verified. To this end, total RNA was extracted from *C. crispus* alga provided by the ‘Centre national de ressources biologiques marine’ (EMBRC France) according to (19) and analyzed by RT-PCR or cRT-PCR (circular RT-PCR) as described in (20) and using oligonucleotides found in Supplementary Table S2.

**Figure 1.**
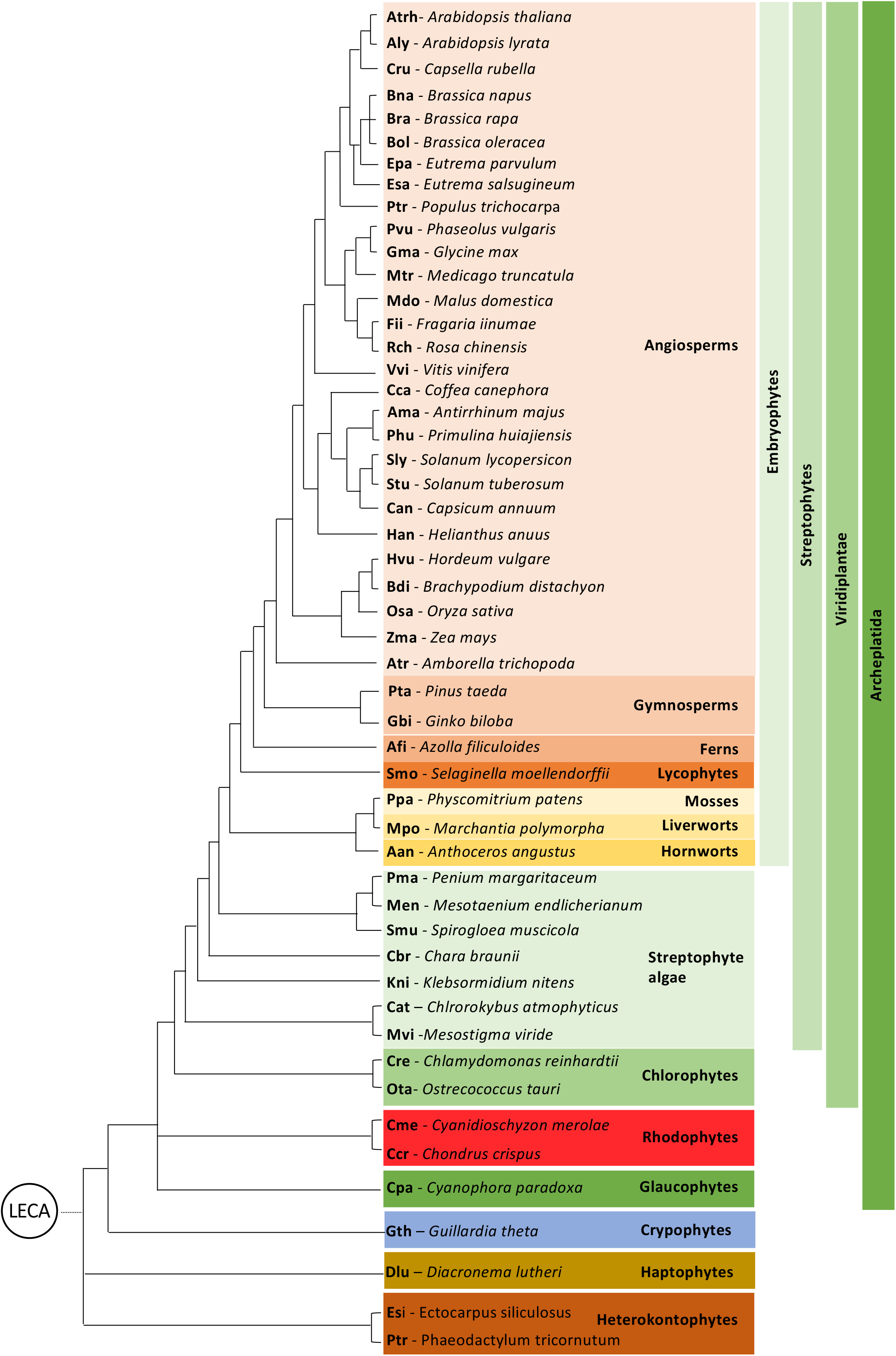
Phylogeny of Archaeplastida with the species included in this study. Two heterokontophytes belonging to the SAR (Stramenopiles, Alveolates, Rhizaria) clade are included. The tree has been constructed from various published studies, in particular (28-31). LECA: Last Eukaryotic Common Ancestor. For each species, a three letters abbreviation is indicated in bold.

We also annotated some of the sequences as tRNA-like sequences. This tRNA-like group comprises true tRNA pseudogenes but likely also highly divergent tRNA molecules which functions remain to be elucidated. Without experimental studies, it is difficult to discriminate between the two categories. In addition, numerous mitochondrial or plastidial tRNA gene sequences inserted into nuclear genomes, and reported so far not be expressed and functional (18), or contaminating the nuclear DNA preparations are recognized as true tRNA genes by tRNAscan-SE. They were annotated and can be downloaded in xls format. Finally, tRNA genes of bacterial origin likely due to DNA contamination were also discarded.

All tRNA sequences, annotations, and biological information are stored in a database implemented in PostgreSQL version 12 (https://www.postgresql.org/). To increase the traceability, we added the genome resources (genome assembly version and source) and the software and versions used for automatic annotation. The web application querying the underlying database is running in a Java-based Apache Tomcat 8.5 environment (https://tomcat.apache.org/). To manage the increasing number of genomes to process and improve the data quality, some automation and systematic data checks are included in a private area of the application.

## RESULTS AND FUTURE DIRECTION

In total, 34403, 1661, and 911 tRNA genes were registered from the nuclear, plastidial, and mitochondrial genomes. Among the 51 photosynthetic organisms, 4 plastidial and 13 mitochondrial genome sequences are not available.

At the level of nuclear genomes analysis, the level of curation was variable (Figure 2). tRNA genes of bacterial origin retrieved on scaffolds (*i.e*. not localized in chromosomes) by tRNAscan-SE 2.0 were deleted: 47 in *Coffea canefora*, 40 in *Mesotaenium endlicherianum*, 35 in *Mesostigma viride*, and 20 in *Chondrus crispus*. Similarly, many plastidial or mitochondrial tRNA genes were detected; their number greatly fluctuates from 0 in the hornwort *Anthoceros angustus* to 1978 in the fern *Azolla filiculoides*. When located on chromosomes, the presence of these tRNA genes is due to organellar DNA insertion into the nuclear genomes; when unplaced (*i.e*. on non-assembled scaffolds), it is difficult to discriminate between organellar DNA insertion and organellar DNA contamination, in particular when genomes have not been assembled into chromosomes. Finally, depending on the quality of the genome assembly and annotation, the number of tRNA-like/tRNA pseudogenes is highly variable. More than 10 000 sequences from the nuclear genomes of the fern *A. filiculoides* or of the gymnosperm *Pinus taeda* belong to this category as well as *circa* 6 000 sequences in the genome of the streptophyte alga, *Chara braunii*. As shown in Figure 2, the remaining true tRNA genes may represent not more than 10% (*e.g*. in *P. taeda* or in *C. braunii*) of the initial sequences retrieved thanks to tRNAScan-SE. The presence of numerous tRNA-like/tRNA pseudogenes may be explained by genome size expansion for example by whole-genome duplication as is the case in ferns or in gymnosperms for instance (21) (22). In gymnosperms, the high proportion of tRNA pseudogenes is in agreement with the numerous pseudogenes observed in conifer genomes and is correlated with the accumulation of transposable elements (23).

**Figure 2.**
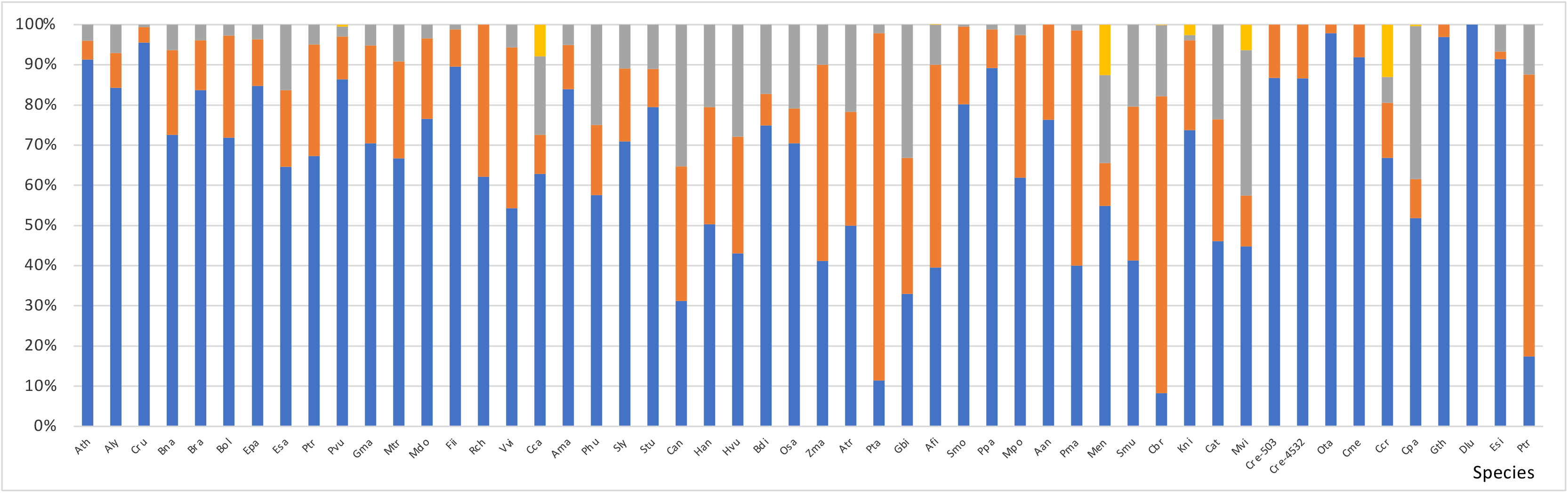
Histogram showing the percentage of nuclear tRNA genes, as estimated from our study. Species are indicated by the three letters code presented in Figure 1. Note that two *Chlamydomonas reinhardtii* strains (Cre-503 and Cre-4532) have been studied. The color code is as followed: true tRNA genes in blue, tRNA-like/tRNA pseudogenes in orange, tRNA genes inserted into the nuclear genome or due to contamination by mitochondrial or plastidial DNA in grey, and tRNA genes corresponding to external bacterial contamination in yellow.

All photosynthetic organisms studied here possess intron-containing nuclear tRNA genes. In embryophytes (Figure 1), around 5% of them possess a classical intronic sequence between positions 37 and 38 and correspond to two tRNA gene families (tRNA^Tyr^ and tRNA^eMet^). This proportion increases up to 13.8% in Brassicaceae, such as *A. thaliana*, where a cluster of tRNA^Tyr^ genes exist (11). In *Ginko biloba*, tRNA^Ile^(TAT) genes also contain classical intronic sequences. In streptophyte algae and chlrorophytes, the number and identity of intron-containing nuclear tRNA genes is more variable and usually higher. The permuted tRNA genes identified in the rhodophyte *Cyanidioschyzon merolae* (24) were also annotated as well as the intronic sequences located at non-canonical positions in the D-, T- or variable loops (25). Similarly, scanning the nuclear genome of another rhodophyte, *Chondrus crispus*, several essential tRNA genes were considered as pseudogenes when using tRNAScan-SE. Manual inspection allowed us to identify introning sequences in either the D-loop between positions 20 and 21, or in the T-loop between positions 58 and 59 of these tRNA genes. Interestingly, the introns in the D- or T-loops are always at the same position and tRNA^Tyr^ and tRNA^eMet^ genes have two introns, the canonical one in the anticodon region plus a second one in either the Dor T- loop (Figure 3). The expression of these *C. crispus* tRNAs and the splicing of their introns were supported by RT-PCR and cRT-PCR experiments conducted for three of them, tRNA^Cys^ (with an intron in the D-loop), tRNA^Trp^ (with an intron in the T-loop), and tRNA^Tyr^ (with two introns in the D- and anticodon loops). In the three cases, correctly spliced tRNAs ending with a CCA triplet were obtained (Supplementary Figure S1).

**Figure 3.**
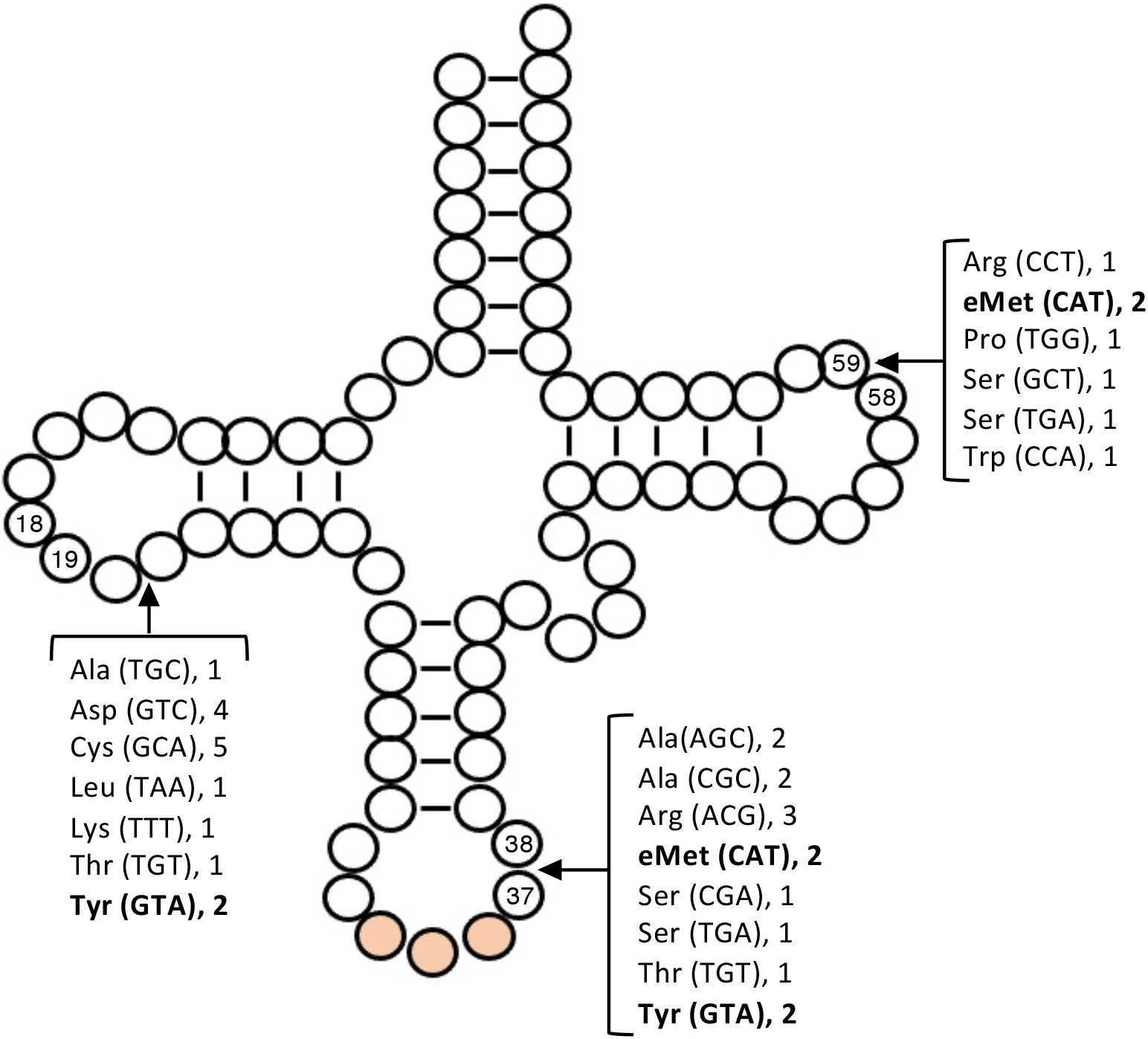
Insertion points introns in *Chondrus crispus* tRNA genes. The typical secondary structure of a mature tRNA is presented. The anticodon is represented with an orange circle. Insertion points of canonical introns in the anticodon loop, and of non-canonical introns in either the D or the T-loops are indicated by arrows. The tRNAs possessing introns are listed by their cognate amino acid, their anticodon in parenthesis, and the number of isodecoders.

Regarding mitochondria-encoded tRNAs, the novel annotated genomes extend our knowledge on the number and identity of this population of tRNA molecules. Interestingly, even only considering embryophytes, there is a great fluctuation in their number: from 0 in the lycophyte *Selaginella moellendorffii* to 126 in the basal angiosperm *Amborella trichopoda*. To compensate for the lack of mitochondria-encoded tRNAs, this likely implies mitochondrial import of a full set of nuclear-encoded tRNAs in the lycophyte, a process experimentally demonstrated in several photosynthetic organisms (26). Conversely, the high number of mitochondria-encoded tRNAs in *A. trichopoda* reflects the presence of numerous horizontal transfers of foreign mitochondrial DNA from green algae, mosses and angiosperms (27). Even with the existence of 126 mitochondria-encoded tRNA genes, the only tRNA^Thr^(GGT) is likely not sufficient to decode the four threonine ACN codons, thus likely requiring the need to import at least one nuclear-encoded tRNA^Thr^. Based on the lack of mitochondria-encoded tRNA species, imported tRNAs were proposed in the PlantRNA database for all sequenced mitochondrial genomes.

Nuclear tRNA genes are transcribed by RNA polymerase III (pol III). The high number of available nuclear tRNA sequences now allow studying in more detail the important characteristics of pol III transcription. In particular, it will be possible to follow how conserved are the TATA-like elements and CAA motifs found upstream of tRNA gene sequences in photosynthetic organisms and whether their presence is linked to plant evolution or adaptation to the environment. Similarly, stretches of T residues normally involved in pol III transcription termination are not always present or are variable in length or number. Whether this variability reflects differences in genome organization and expression remains to be elucidated.

In PlantRNA 2.0, no emphasis has been given on related biological information such as aminoacyl-tRNA synthetases or modification enzymes and work needs to be done to fill this gap. From an informatics point of view, many steps required to implement the PlantRNA database have now been automatized. The main bottleneck in updating more regularly the database remains the manual curation. We now plan, with the large number of curated tRNA sequences, to implement a neural network so that any new genome is accurately annotated and the data quickly incorporated into the database. Finally, the incorporation into the database of tRNA and tDR expression profiles as well as the modified nucleotide composition of each tRNA species is biologically relevant but will first require more experimental work.

## DATABASE AVAILABILITY

PlantRNA is accessible freely at http://PlantRNA.ibmp.cnrs.fr. All published data performed with the help of the PlantRNA database should refer to this article.

## SUPPLEMENTARY DATA

Supplementary data are available at NAR online.

## ACKNOWLEDGEMENTS

We would like to thank Marie Bernhard and Thalia Salinas-Giegé for technical support, and Alexandre Berr for critical discussion.

## AUTHOR CONTRIBUTIONS

LD conceived the project, curated manually the data, and wrote the manuscript. VC developed the workflow to annotate tRNA, processed all the genomes, and contributed to the design of the database and the data integration. GP implemented the PlantRNA database and the web interface and DP adapted the tRNAscan import R package to split the tRNA sequences. All authors contributed to the final manuscript.

