## Supplementary material for "PlantRNA 2.0: an updated database dedicated to tRNAs of photosynthetic eukaryotes": Figure S1

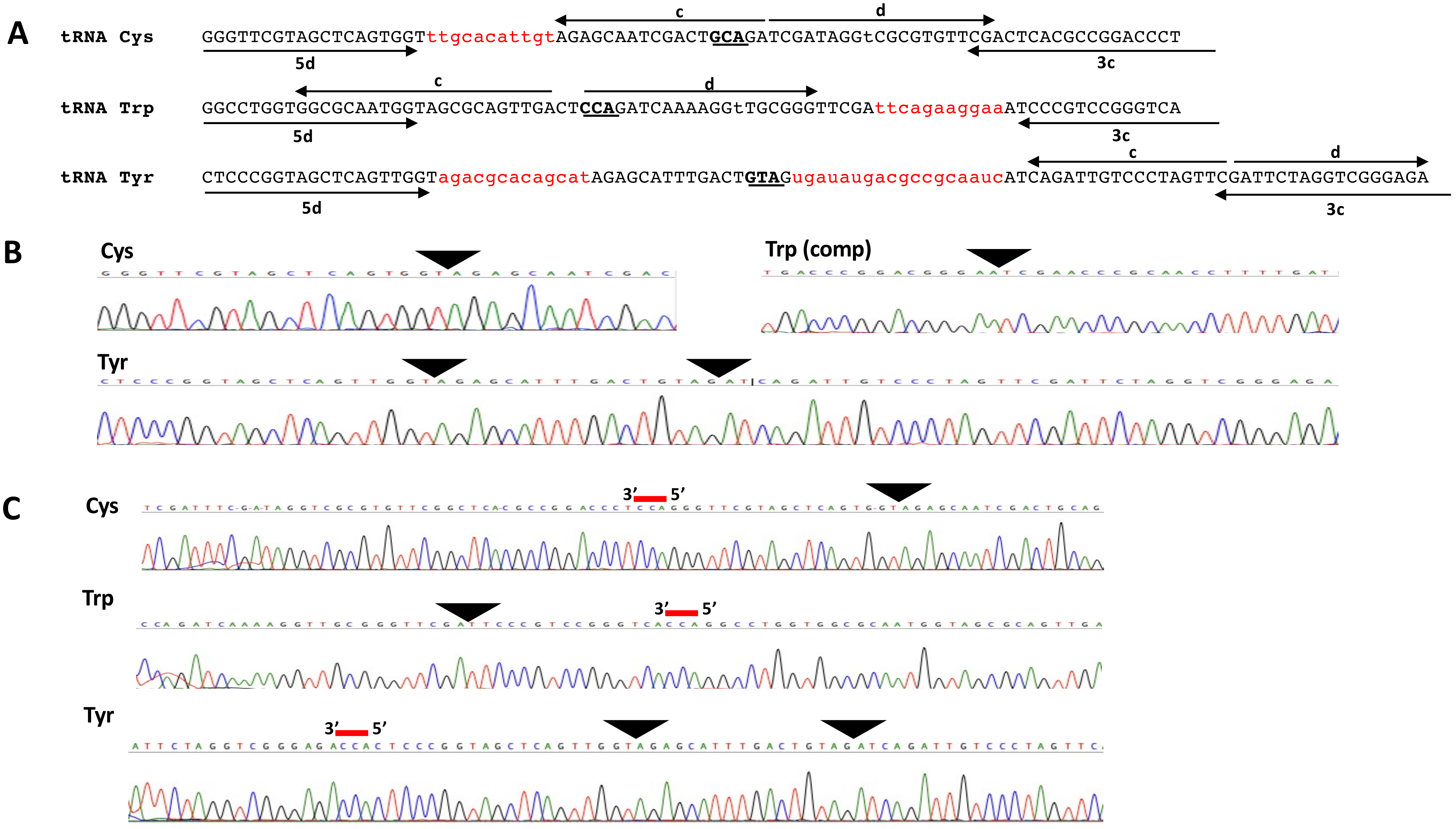


**Figure S1**: **A)** Sequences of nuclear-encoded tRNACys, tRNATrp and tRNATyr genes from *Chondrus crispus*. Sequences indicated by arrows were used as primers for either RT-PCR (5d, 5c) or cRT-PCR (d, c) experiments. Putative introns are in red. Anticodons are in bold and underlined, **B)** and **C)** Mapping of the mature tRNA extremities by RT-PCR and cRT-PCR. Sample sequence of one of the clones obtained from amplification products. Post-transcriptionally added CCA triplets are indicated by a red bar. Black arrow heads indicate the position of spliced introns.
